# Multi-omics analysis reveals cross-organism interactions in coral holobiont

**DOI:** 10.1101/2021.10.25.465660

**Authors:** Toru Maruyama, Michihiro Ito, Satoshi Wakaoji, Yusuke Okubo, Keigo Ide, Yohei Nishikawa, Hiroyuki Fujimura, Shoichiro Suda, Yoshikatsu Nakano, Noriyuki Satoh, Chuya Shinzato, Kei Yura, Haruko Takeyama

## Abstract

Corals create an ecosystem, called a holobiont, with intracellular algae (zooxanthellae) and resident bacteria. Zooxanthellae and some bacteria play major roles in the physiological properties of the coral host. However, because of the difficulties in experimental verification of cross-organism interactions, the mechanisms underpinning these interactions are largely unknown. To address this, we here generated and then analyzed multi-omics datasets for corals, zooxanthellae, and bacteria collected at Okinawa, Japan, from November 2014 to September 2016. Using cross-organism co-expression analysis, we successfully characterized the host–alga relationship in the coral holobiont. Specifically, we observed that the coral host dominates the zooxanthellae. The multi-omics analysis also suggested that infection with coral-associated bacteria *Endozoicomonas* likely involves coral-like ephrin ligands, triggering an immune response of the coral host. This study highlights the potential of the multi-omics approach to elucidate coral–microbe interactions.

## Introduction

Corals form an ecosystem, called a holobiont, with intracellular dinoflagellates and resident microbiota. The intracellular dinoflagellates, zooxanthellae, play critical roles in the coral host, as they are important photosynthate providers. For instance, zooxanthellae produce more than 90% of metabolites required for coral respiration, via photosynthesis ^1,2^. These dinoflagellates also synthesize other beneficial substances, such as mycosporine-like amino acids ^3,4^. Collapse of the symbiotic relationship is detrimental to the coral and has been a major factor in the worldwide decline in coral populations ^5^. The breakdown of symbiosis, known as coral bleaching, is triggered by stress factors such as rising sea temperature; however, the mechanism underlying this phenomenon is far from being elucidated ^6^. Further, while the above symbiotic relationship is generally considered mutually beneficial ^1,2^, some controversy surrounds this notion, and some researchers hypothesize that the coral host exerts a dominant control over the intracellular symbiont ^7^.

A close relationship between the coral and bacteria is also expected, as first proposed in coral probiotic hypothesis ^8^. It has been reported that the bacterial community structure changes in response to the conditions to which the coral host is subjected, including bleaching ^9^, disease ^10–13^, and temperature ^14^. Further, alteration of the bacterial flora in coral mucus by antibiotics causes coral bleaching and death^15^. The coral microbiome is completely different from that of the surrounding seawater ^16^ and consists of commensal bacteria ^17,18^. Bacteria from the *Endozoicomonas* genus have been identified in coral microbiomes worldwide and have been proposed to reside inside coral tissues and cells ^18,19^. Further, it has been reported that the health of the coral host deteriorates when the *Endozoicomonas* population declines, suggesting that these microorganisms exert a beneficial effect on the coral host ^9,13,15^. However, *Endozoicomonas* have been suggested to be pathogenic in fish ^20,21^, and the mechanisms by which they contribute to coral health are largely unknown.

Although the relationship between the intracellular algae and resident bacteria is important for corals, the underlying molecular mechanisms have not been fully elucidated because of the difficulties associated with the experimental investigation of this complex biological system. Many intracellular algae and resident bacteria are unculturable ^22,23^, which hampers the attempts to set up an experimental model of symbiotic interactions. Consequently, an alternative, data-driven approach involving simultaneous omics profiling of the host and microbe, is used to understand such cross-organism interactions ^24^. In coral research, simultaneous profiling of coral and zooxanthellae mRNA revealed a repertoire of genes in both organisms ^25^. In the current study, we have extended the multi-omics approach to decipher interspecific crosstalk in the coral holobiont. We profiled the (i) coral and (ii) zooxanthellae transcriptome; (iii) 16S rRNA-based coral microbiome; and (iv) bacterial genome. Integrative analyses of the multi-omics datasets successfully highlighted complex interactions among corals, zooxanthellae and coral microbiome.

## Results

### Multi-omics profiling of coral holobiont

We collected the scleractinian coral *Acropora tenuis* at three sampling sites, Ishikawabaru (*Is*), Kyuusanbashi (*Ky*) and Sesoko-minami (*Mi*) around Sesoko Island, Okinawa, Japan (Supplementary Fig. 1), between November 2014 to September 2016. *Is* is located in a reef lagoon and is close to a sewage plant and a tuna farm. *Mi* and *Ky* are in open areas with low impact from human activity. We collected 63 samples from seven coral colonies (Supplementary Table 1). The corals showed signs of bleaching in August and September 2016 but appeared healthy at other sampling times (Supplementary Fig. 2).

We simultaneously generated multi-omics data of coral, zooxanthella, and bacterium for the collected samples (Fig. 1). Briefly, we extracted mRNAs of coral and zooxanthella from the total RNA of the holobiont and sequenced them simultaneously. We aligned the mRNA-seq sequence reads to the genome of *A. tenuis* ^26^ to determine the mRNA profiles of the coral hosts. For the dataset, 33.6–69.0% (average 52.6%) of the reads were aligned to *A. tenuis* genome (Supplementary Table 2). We then extracted the unmapped sequence reads for *de novo* assembly to obtain zooxanthella transcripts. Finally, we analyzed the obtained 325,653 contigs by similarity searching against publicly available transcript sequences of corals and zooxanthellae ^4,26–29^. Accordingly, 106,269 and 59,265 contigs shared similarity with the coral and zooxanthella sequences, respectively (Supplementary Fig. 3a), and were characterized by distinct GC content distributions (Supplementary Fig. 3b). The unclassified contigs were largely short (Supplementary Fig. 3c) and had a GC content close to that of corals (Supplementary Fig. 3b). Therefore, we concluded that the majority of the zooxanthella transcripts were successfully extracted using the described approach. We re-aligned the unmapped reads to the zooxanthella transcripts and determined the gene expression profiles of the intracellular algae. We also amplified the 16S rRNA V1–V2 region from the total holobiont RNA by RT-PCR and sequenced the resultant amplicons to obtain information on coral-associated bacterial communities. Finally, we analyzed the genome sequence of *Endozoicomonas* sp. ISHI1, isolated from *A. tenuis* at the *Is* site.

**Fig. 1.**
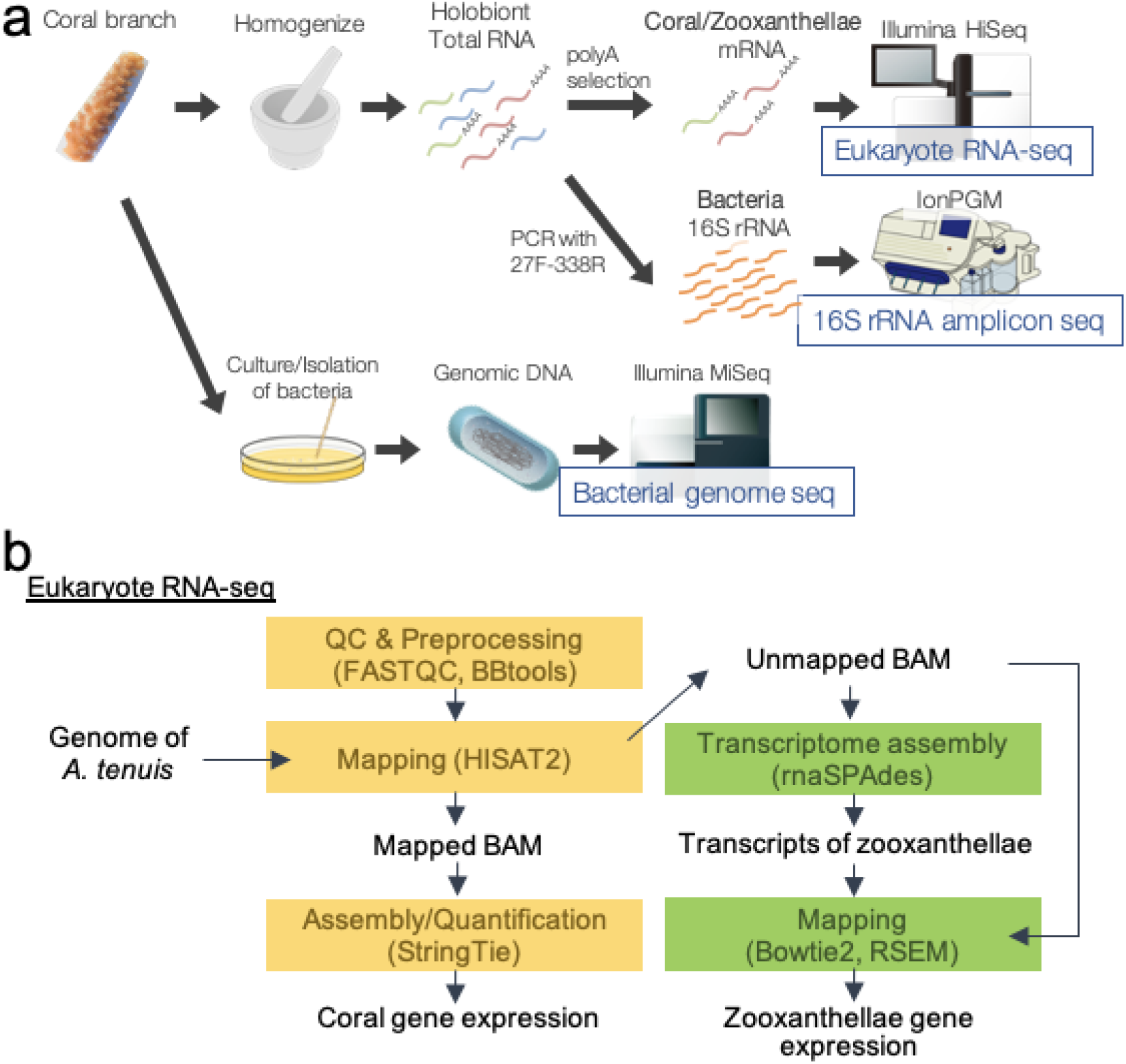
Multi-omics profiling of the coral holobiont. **a** Overview of experimental procedures for acquiring multi-omics datasets. **b** Bioinformatics pipeline for multi- omics analysis of the coral holobiont.

### Season and colony-dependence of coral gene expression

We analyzed the coral transcriptome using weighted gene co-expression network analysis (WGCNA) and identified 36 co-expression gene sets, henceforth referred to as “modules”. Each module showed a different expression pattern, and their sizes ranged from 55 to 321 genes. Twenty modules showed seasonal variation, whereas the other 16 were differentially expressed in specific colonies (Fig. 2a,b). For example, the “green” module showed season-dependent expression and was highly expressed during summer (2015/08, 2016/08, and 2016/09) (Fig. 2a), while the “darkgrey” module was specifically expressed in Is3 colony regardless of the season (Fig. 2b).

**Fig. 2.**
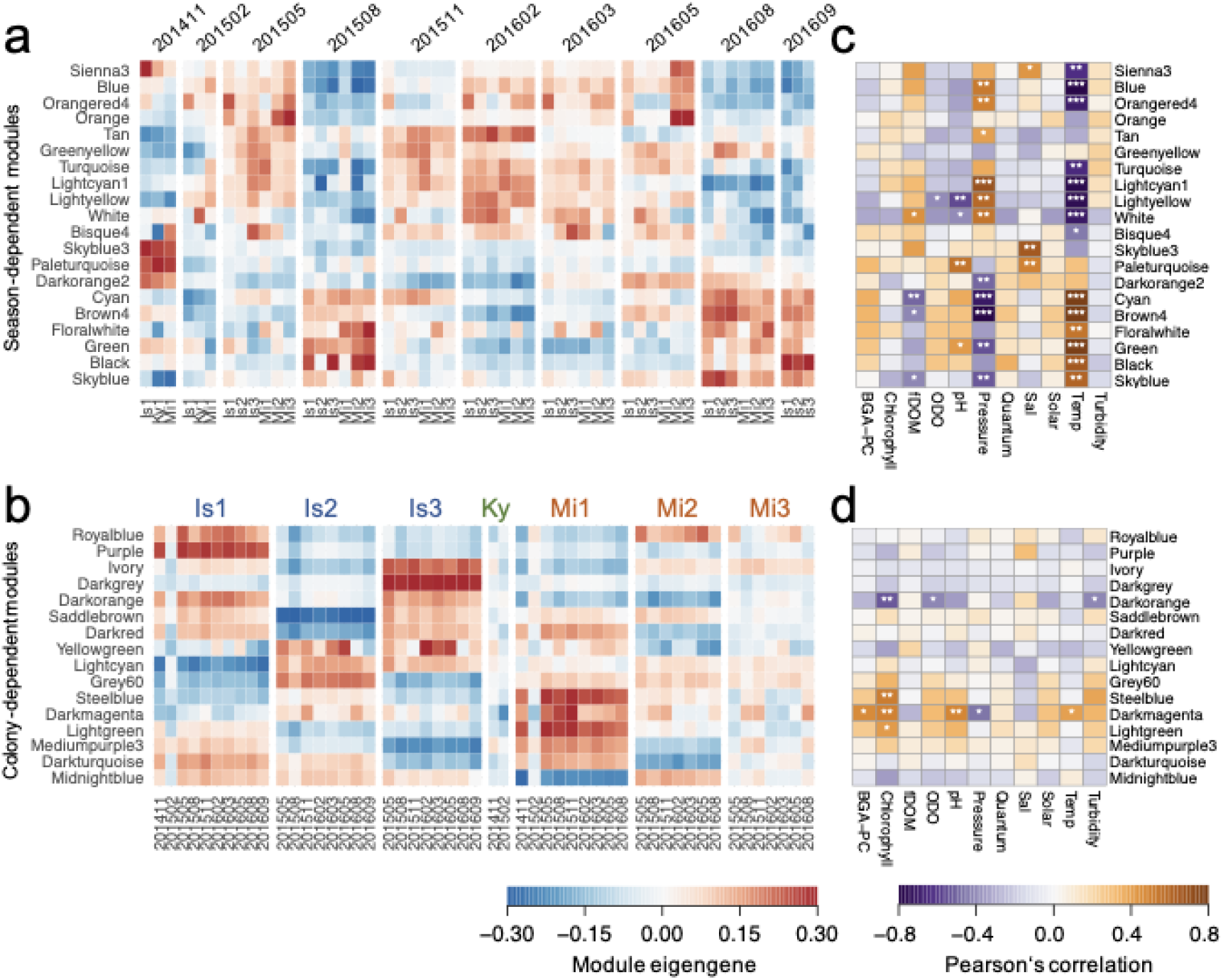
Season- and colony-dependent patterns of coral gene expression. **a,b** Module eigengenes (the expression level of modules) of season-dependent modules (**a**) and colony-dependent modules (**b**). **c,d** Correlation between environmental parameters and module eigengenes. Statistical significance of the correlation is denoted as follows: * p < 0.05, ** p < 0.01, *** p < 0.00001.

Among the 20 season-dependent modules, 18 showed a significant correlation (p < 0.05) with at least one environmental parameter (Fig. 2c). The expression of 15 modules was significantly correlated (or anti-correlated) with water temperature. Among them, the positively correlated modules were significantly associated with GPCR signaling (black, cyan, floralwhite, and green), organization of the extracellular matrix and bone (skyblue), ion channel activity (cyan), and the exosome/phagosome (brown4 and skyblue) (Supplementary Fig. 4a, and Supplementary Tables 3 and 4). The negatively correlated modules were significantly associated with protein/peptide degradation process (lightyellow, turquoise, and white), cell replication (blue and sienna3), and sphingolipid metabolism (orangered4) (Supplementary Fig. 4b, and Supplementary Tables 3 and 4). Two modules (skyblue3 and paleturquoise) were correlated with salinity and pH; however, no significant functional enrichments were apparent. The “orange” module was significantly associated with spermatogenesis (Supplementary Table 3). Further, this module was upregulated in the spawning season (May 2015 and 2016), suggesting that it was indeed associated with the reproduction process.

Sixteen colony-dependent modules were not as well annotated as the season-dependent modules, as determined by similarity searches against Swiss-Prot and KEGG databases (Supplementary Fig. 5). Seven colony-dependent modules were significantly associated with the process of DNA integration, recombination, and transposition (royalblue, saddlebrown, darkred, lightcyan, steelblue, darkturquoise, and midnightblue) (Supplementary Fig. 6). These observations suggest that the modules involved in intraspecific variations were enriched with functionally unknown genes, with some of them acquired via genetic recombination and transposition.

### Seasonal changes in zooxanthella gene expression

We also applied WGCNA to the zooxanthella transcriptome and identified 27 modules. Among them, 22 modules showed seasonal variation, whereas the remaining five were differentially expressed in specific colonies (Fig. 3a,b). To evaluate how changes in zooxanthella populations impacted the analysis, we examined the origin of zooxanthella clades that each module was derived from. More than 75% of all transcripts shared the highest similarity with the sequences of Clade C zooxanthellae (*Cladocopium* sp. clade C or *Cladocopium goreaui*) ^30^, and every module exhibited a similar trend (Supplementary Fig. 7). This suggests that *Cladocopium* is the dominant zooxanthella type regardless of the season and colony and that the modules reflect transcriptional changes to a greater extent than zooxanthella population changes.

**Fig. 3.**
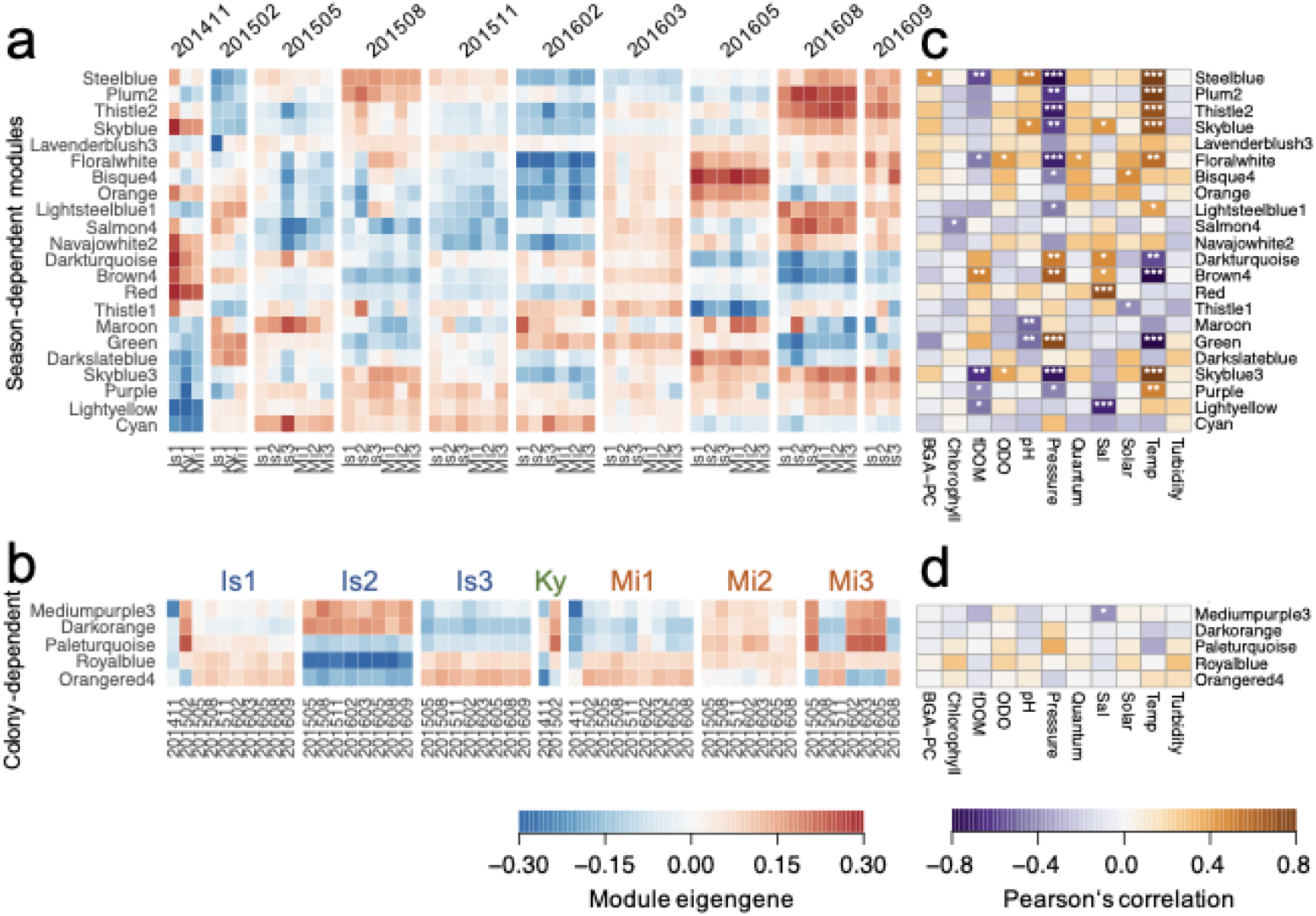
Season- and colony-dependent patterns of zooxanthella gene expression. **a,b** Module eigengenes of season-dependent modules (**a**) and colony- dependent modules (**b**). **c,d** Correlation between environmental parameters and module eigengenes. Statistical significance of the correlation is denoted as follows: * p < 0.05, ** p < 0.01, *** p < 0.00001.

Among the 22 season-dependent modules, 18 were significantly correlated (p < 0.05) with at least one environmental parameter (Fig. 3c). Eleven modules were significantly correlated (or anti-correlated) with water temperature, with eight modules positively correlated with water temperature. Of these eight modules, the “floralwhite” and “lightsteelblue1” modules were significantly associated with cilium movement; “plum2” and “skyblue” were significantly associated with photosynthesis; “skyblue3” was significantly associated with cell cycle; and “thistle2” was significantly associated with superoxide regulation (Supplementary Tables 5 and 6). In addition, the “brown4” module, which was negatively correlated with water temperature, was associated with the interleukin 17 signaling pathway (Supplementary Table 6). Finally, the “red” module was significantly associated with carbon fixation and nitrogen metabolism and was highly correlated with salinity (Supplementary Tables 5 and 6).

### Cross-species co-expression network analysis

To determine the factors affecting the coral and zooxanthella transcriptomes, we calculated the correlations for their gene expression profiles. The gene expression profiles of the corals were similar among samples originating from the same colony (Supplementary Fig. 8a,d), while the profiles of zooxanthellae were similar among samples collected at the same sampling time (Supplementary Fig. 8b,e). These observations suggest that many genes are independently regulated in the coral hosts and in zooxanthellae.

We then investigated cross-species gene co-expression in the corals and zooxanthellae to estimate mechanisms underlying the coral–algal symbiosis. Specifically, to understand mechanisms of their interaction underpinning coral bleaching, we selected the modules that were significantly correlated with zooxanthella cell density (Supplementary Fig. 2) and calculated the co-expression among them (Fig. 4). Zooxanthella cell densities were positively correlated with coral modules associated with DNA replication (sienna3), cell cycle (blue), and sphingolipid biosynthesis (orangered4). The coral modules co-expressed with zooxanthella modules were associated with carbon fixation and nitrogen metabolism (red). However, the carbon fixation module (red) showed negative correlation with zooxanthella module associated with the DNA replication process (skyblue3). These findings suggest that carbon fixation by zooxanthellae contributes to the growth of the coral host, but it is not strongly linked to the growth of zooxanthellae themselves.

**Fig. 4.**
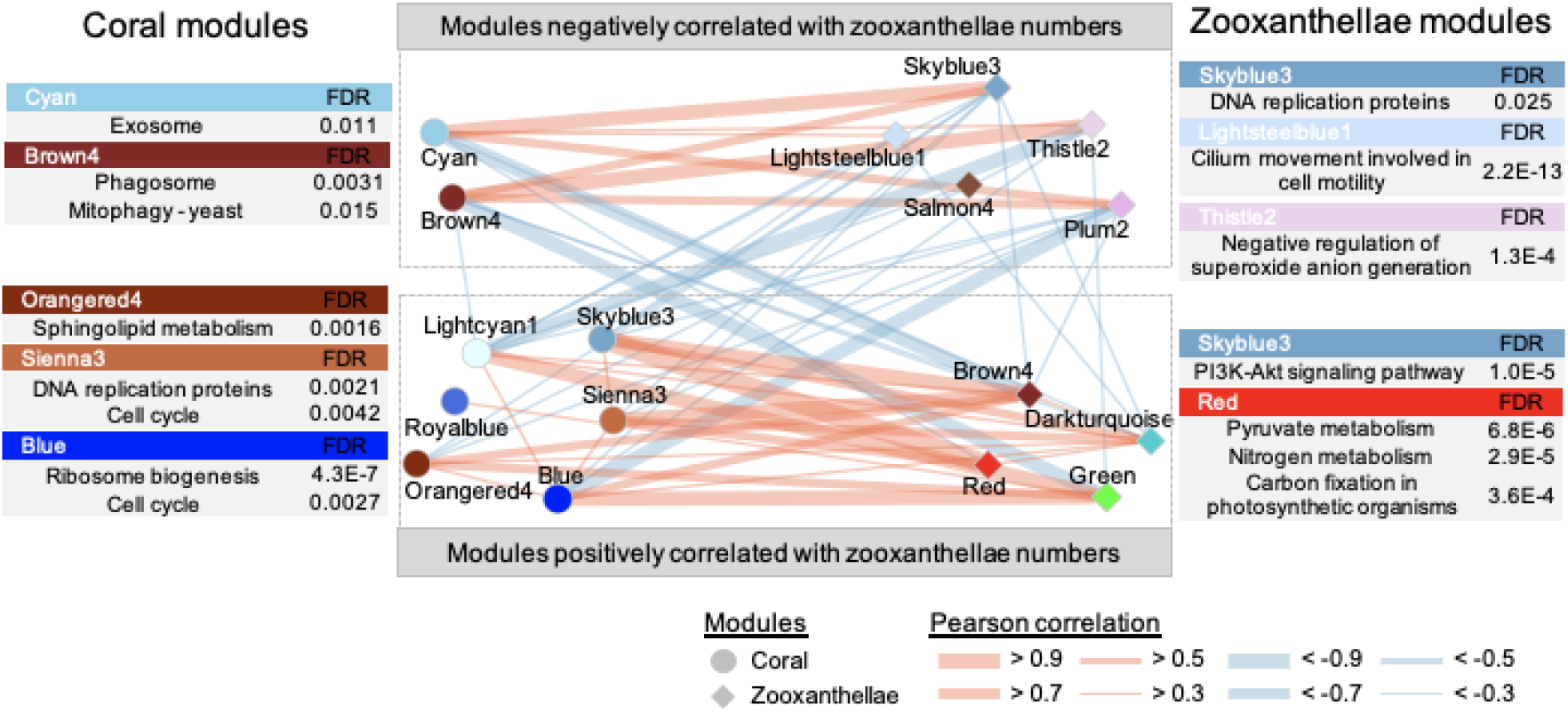
Cross-species co-expression network of the crosstalk between coral and symbiotic algae. Co-expression network created using the WGCNA modules. Circles and diamonds correspond to the modules of coral and symbiotic algae, respectively. Orange and blue edges represent positive and negative correlations of the module eigengenes, respectively.

Zooxanthella cell densities were negatively correlated with coral modules associated with phagosomes (brown4), exosomes (byan), and zooxanthella modules associated with DNA replication (skyblue3), cilium movement (lightsteelbule1), and regulation of reactive oxygen species generation (thistle2). These observations suggest that zooxanthella cell density is actively regulated by the coral host via phagocytosis and exocytosis and that the expelled zooxanthellae may acquire motile behavior by retaining the flagella, as previously suggested ^2^.

### Site and colony-dependent community structure of coral microbiome

We analyzed the bacterial composition of coral microbiome and the surrounding seawater using total RNA-based 16S rRNA amplicon sequencing. The coral microbiome was mainly composed of microorganisms from the *Rickettsiales* family (*Anaplasma-like*) ^31^, *Endozoicomonas*, and *Cyanobacteria* Group VIII (Fig. 5a). Further, it was dominated by 10 operational taxonomic units (OTUs) (Fig. 5b), which accounted for >60% of the coral microbiome, with <0.3% detected in the surrounding seawater. Principal coordinate analysis (PCoA) using unweighted UniFrac distance revealed that the data points for microbiomes from corals and seawater grouped separately (Fig. 5c).

**Fig. 5.**
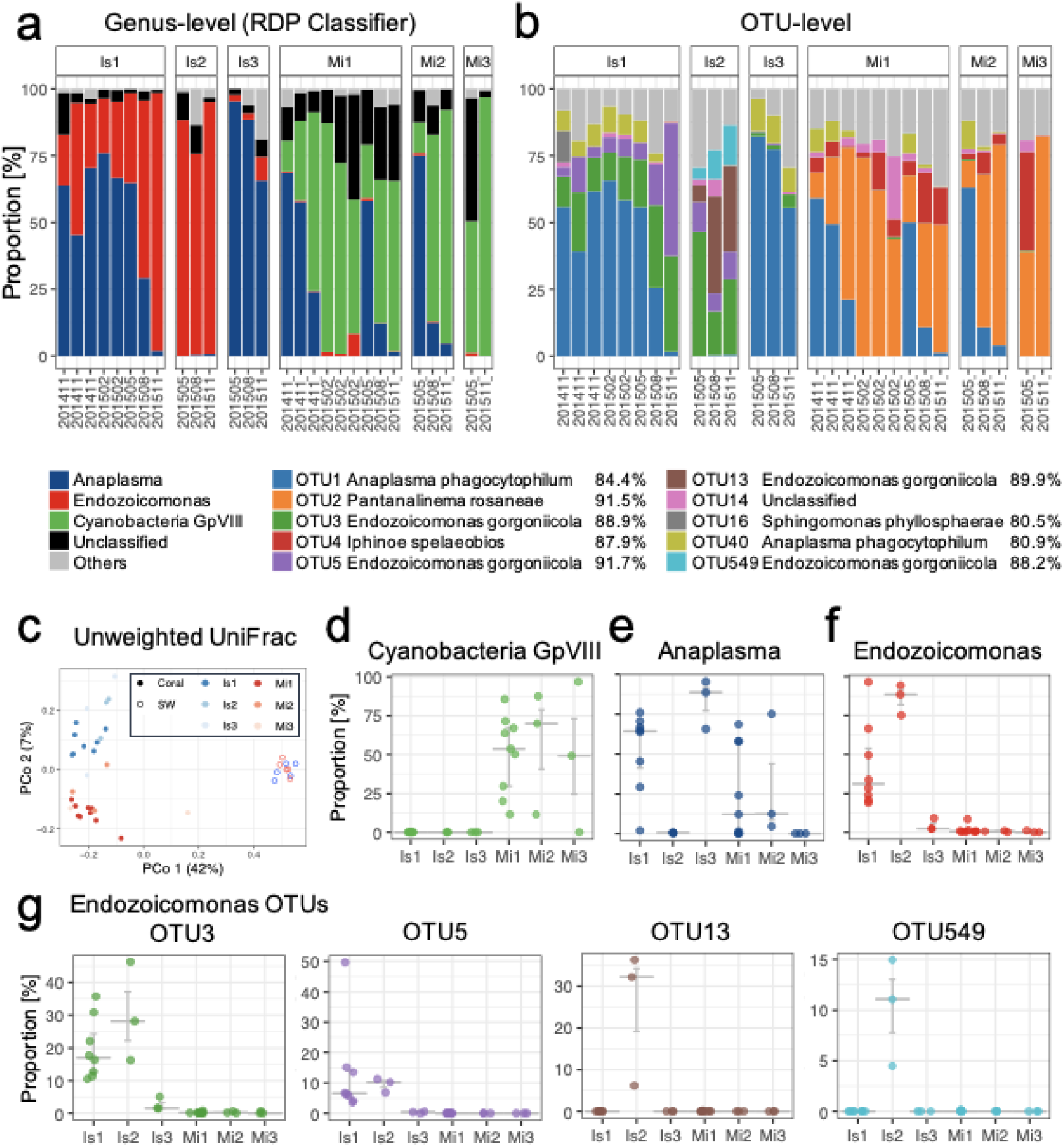
Composition of the coral microbiome. **a,b** Composition of the coral microbiome at the genus level (**a**) and OTU level (**b**). Genera and OTUs with an abundance of less than 10% in all samples are denoted as “Others”. OTU genera were based on top hits after similarity search against the SILVA database. **c** Embedding of the community structure by principal coordinate analysis (PCoA) using unweighted UniFrac distance. **d–f** Proportions of *Cyanobacteria* GpVIII, *Anaplasma*, and *Endozoicomonas* in each colony. **g** Proportions of four major *Endozoicomonas* OTUs.

Some genera and OTUs were specific to the sampling site or the colony. For instance, we only detected *Cyanobacteria* Group VIII at the *Mi* site (Fig. 5d). By contrast, we identified *Anaplasma-*like bacteria, which was reported as a marker of coral disease susceptibility ^32^, at *Is* and *Mi* sites, except for the Is2 and Mi3 sites (Fig. 5e). Finally, we detected *Endozoicomonas*, a genus recognized as a commensal of diverse marine organisms, including corals at several geographic locations, mostly at the *Is* site, especially the Is1 and Is2 colonies (Fig. 5f). Distribution patterns of the species from the genus *Endozoicomonas* differed. We typically identified OTU3 and OTU5 at Is1 and Is2, whereas OTU13 and OTU549 were specific to Is2 (Fig. 5g). Further, the UniFrac distance analysis indicated the highest similarity among samples originating from the same colony or the same sampling site (p < 0.01, PERMANOVA) (Supplementary Fig. 8c,f). In contrast with the gene expression profiles of corals and symbiotic algae, the composition of coral microbiome was strongly affected by the sampling site (Supplementary Fig. 8). This suggests that the microbiomes are regulated not only by the coral host but also by other factors that do not directly influence coral gene expression.

### Associations between coral microbiome and the coral host

We only detected *Anaplasma-*like and *Endozoicomonas* bacteria in specific coral colonies (Fig. 5e,f). We therefore examined the genes differentially expressed in these colonies to determine the impact of each genus on the coral host. The analysis revealed that 1,500 and 1,425 genes were up or downregulated, respectively, in the colonies harboring *Anaplasma*-like bacteria (Is1, Is3, Mi1, and Mi2) (Supplementary Table 7). Further, 1,690 and 1,984 genes were up or downregulated in the colonies associated with *Endozoicomonas* (Is1 and Is2) (Supplementary Table 8). Gene sets related to immune reaction (e.g., positive regulation of interleukin 1 or regulation of killing of cells of other organisms) and vesicle formation (e.g., vesicles budding from membrane or phagolysosome assembly) were significantly enriched in the upregulated genes in colonies harboring *Endozoicomonas* (Supplementary Table 9). The upregulated genes included several P2X purinoceptor 7 (P2RX7) genes (Gene ID: MSTRG.12568 and MSTRG.9878) (Fig. 6a, Supplementary Table 9). P2RX7 is associated with host–pathogen interactions, such as elimination of intracellular parasites ^33^, and is conserved in cnidarian species ^34^. We speculated that *Endozoicomonas* triggers inflammatory reactions by activating host P2RX7. The upregulated genes also included ephrin receptor genes (Gene ID: MSTRG.5097 and MSTRG.31636) (Fig. 6b, Supplementary Table 8). Ephrin ligands are specific to eukaryotes; however, their genes have also been identified in the genomes of coral-associated bacteria, such as *Endozoicomonas montiporae* ^35^. The ligands are thought to be involved in the infection process since they mediate several cellular functions, including endocytosis ^35^. These observations suggest that the ephrin ligand is likely involved in *Endozoicomonas* infection.

**Fig. 6.**
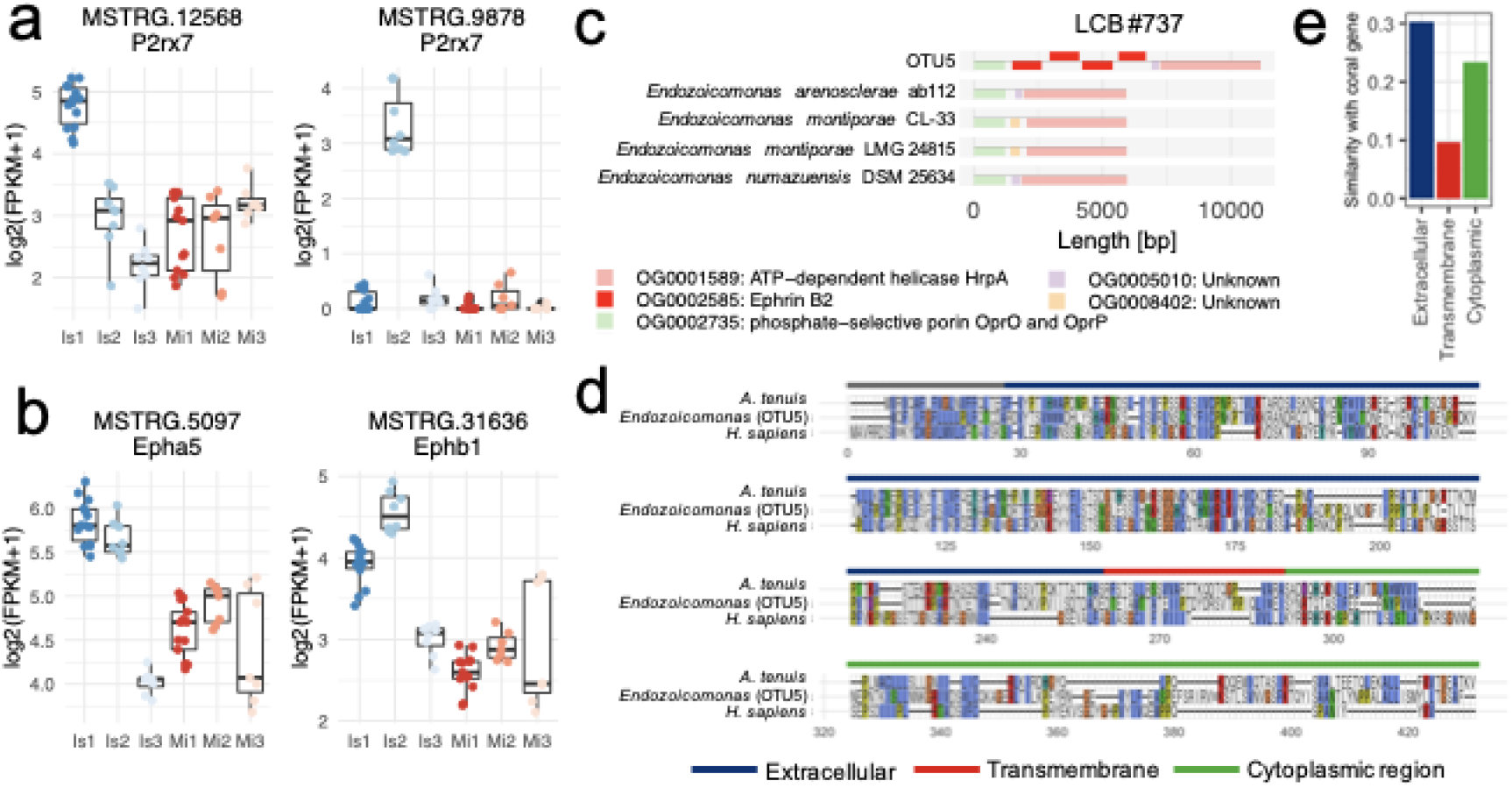
Association between coral gene expression and the microbiome. **a,b** Genes differentially expressed in colonies harboring *Endozoicomonas* (Is1, Is2): *P2RX7*(**a**) and ephrin ligand genes (**b**). **c** Locally collinear block (LCB) around ephrin-B2 genes in OTU5. Genes are colored by orthogroups defined with Orthofinder. **d** Multiple alignment of ephrin ligand genes in coral (*A. tenuis*), *Endozoicomonas* (OTU5), and human. **e** Similarity between the ephrin genes of OTU5 and *A. tenuis*.

We then successfully isolated *Endozoicomonas* species (*Endozoicomonas* sp. ISHI1) corresponding to OTU5 and analyzed its genome sequence. Subsequently, we identified eight ephrin ligand genes (e.g., NGGDMNLC_01928) (Supplementary Table 10) and found that their genomic locations were different from those in *E. montiporae* (Fig. 6d). These findings indicate that OTU5 acquired the host-like ephrin ligand genes at a different time than *E. montiporae* and that the ephrin ligand has an evolutionarily advantageous role for coral–bacteria interactions.

We next performed a similarity search for the eight newly identified ephrin ligand genes using the NCBI nr database. The top hits were ephrin ligand genes from *Acropora* corals. The extracellular regions of the coral and *Endozoicomonas* ephrin ligand genes shared higher similarity (30.4% identity) than those of the transmembrane regions (9.7% identity) and cytoplasmic regions (23.3% identity) (Fig. 6e). Hence, the extracellular domain might be important for establishment of the relationship between *Endozoicomonas* and the coral host.

## Discussion

The knowledge of the interspecific interactions within a coral holobiont is limited because of the limitations of experimental methods that are currently available to investigate these interactions. Therefore, in the current study, we used a multi-omics approach to investigate the interactions between coral, zooxanthellae, and resident bacteria. We designed an efficient analysis pipeline and succeeded in determining the coral and zooxanthella gene expression profiles from data obtained by simultaneous sequencing of their transcripts. Then, using a systems biology approach, we successfully determined specific cross-species co-expression patterns between the coral host and zooxanthellae. We also described the inter-colony heterogeneity of the coral microbiome using 16S rRNA amplicon sequencing. Integrated analysis of the coral transcriptome and microbiome revealed gene expression changes that correlated with the existence of coral-associated bacterial species and highlighted the ephrin genes as candidates involved in the coral– bacterium crosstalk. This study demonstrates the potential of a data-driven approach for the analysis of cross-species interactions.

In the current study, the cross-species co-expression network analysis revealed that carbon fixation in zooxanthellae was positively correlated with coral host replication and negatively correlated with zooxanthella replication. This is consistent with previous reports suggesting that the doubling time of zooxanthellae during symbiosis is lower than that of zooxanthellae in culture ^7^. The analysis also revealed that the replication process of zooxanthellae was negatively correlated with the phagocytosis and exocytosis of the coral host. Although several mechanisms of zooxanthella loss have been suggested, it is unclear which of these actually occur in nature ^6^. The results presented herein suggest that the coral host actively regulates the number of intracellular algae by exocytosis-mediated expulsion and by degradation via phagocytosis. While the relationship between the coral and zooxanthellae has long been recognized as mutually beneficial, previous studies have challenged this notion and suggested that the relationship is dominated by coral host ^7^. Similarly, the data-driven approach used in the current study revealed that fixed carbon is mainly used by the coral host, that the zooxanthella replication is suppressed during symbiosis, that zooxanthella density is actively maintained by the coral host, thus suggesting that the coral host presents a dominant interaction in the holobiont.

We also evaluated the inter-colony heterogeneity of coral microbiomes. *Endozoicomonas* dominated the microbiome of two colonies, Is1 and Is2, while they were much less abundant in other colonies. *Endozoicomonas* have been predicted to have an intimate relationship with coral ^18^, but their roles are mostly unknown and controversial ^13,17^. In the current study, we showed that inflammation-related genes were upregulated in corals harboring *Endozoicomonas* species. The upregulated genes contained those involved in defense against bacterial infections, such as *P2X7*. Previous studies have reported that *Endozoicomonas* species are pathogenic to the fish host ^20,21^. In addition, *Endozoicomonas* possess several eukaryotic-like genes like the E3 ubiquitin ligase gene, which have been identified in pathogenic bacteria such as *Salmonella* and *Legionella* ^36^. Although a beneficial effect of *Endozoicomonas* has been previously proposed ^9,15^, the results presented herein suggest that this is questionable, as the coral host exhibited a defense response against *Endozoicomonas*. Furthermore, the bleaching of colonies harboring *Endozoicomonas*, Is1 and Is2, in the summer of 2016 was more severe than that of others colonies. Further verification of the beneficial effects of *Endozoicomonas* species for the coral host is necessary to resolve this.

While we have demonstrated here the potential of data-driven approaches, we acknowledge several limitations of this method. First, the presented analysis revealed that most zooxanthella transcripts were similar to those of *Cladocopium*, suggesting that *Cladocopium* was dominant in the sampled corals regardless of the sampling time and the coral host. However, we were unable to evaluate how the change in zooxanthella composition affected the results of the co-expression network analysis. Regarding this aspect of the current study, we believe that the approach will become more accurate as more zooxanthella genomes become available. Second, the presented analysis did not fully support the hypothesis that *Endozoicomonas* are beneficial to the coral host. However, we believe that further investigation will be necessary to confirm the presented results since we only detected the presence of *Endozoicomonas* in two of the colonies studied, with differences in the *Endozoicomonas* occurrence among colonies at the same sampling site. Using a large number of colonies from the same sampling site might be an effective strategy to implement in future studies.

In conclusion, we here proposed a data-driven approach for the investigation of cross-organism interactions involving coral, zooxanthellae, and bacteria. This approach is potentially applicable to diverse questions in coral biology. For example, it could be used to elucidate the association between bacterial infection and coral disease and the mechanism(s) of resistance to coral bleaching.

## Methods

### Sample collection

Coral branches were collected at three sampling sites [Ishikawabaru (*Is*), Kyusanbashi (*Ky*), and Sesoko-minami (*Mi*)] around Sesoko Island, Japan, from November 2014 to September 2016, by scuba diving (Supplementary Fig. 1). Immediately after collection, the branches were soaked in RNAlater (Ambion, Austin, TX, USA), snap-frozen on ethanol with dry ice on the sampling ship, and stored at – 80 °C until RNA extraction. Before analysis, the branches were crushed and homogenized using an iron mortar and pestle, with the samples soaked in liquid nitrogen. Total RNA was extracted by using the RNeasy RNA extraction kit (QIAGEN, Hilden, Germany).

### Acquisition of environmental data

Environmental parameters, including water temperature (Temp), pH, turbidity, salinity (Sal), solar intensity (Solar), photon flux (Quantum), water depth (Pressure), dissolved oxygen (ODO), fluorescent dissolved organic matter (fDOM), chlorophyll concentration (Chlorophyll), and phycocyanin of blue-green alge (BGA-PC), were measured at the sampling sites using YSI EXO2 water quality sonde (YSI Inc, Yellow Springs, OH, USA) and COMPACT-LW photon sensors (JFE Advantech, Nishinomiya, Japan)

### Coral and zooxanthella transcriptome profiling

Coral and zooxanthella mRNAs were obtained from total RNA, and sequencing libraries were prepared using TruSeq RNA Library Prep kit (Illumina, San Diego, CA, USA). The libraries were sequenced using the Illumina HiSeq 2000 platform (Illumina) in 100-bp paired-end mode. Low-quality reads were removed and adapters trimmed using bbduk in BBTools (sourceforge.net/projects/bbmap/) with the options “ktrim=r k=23 mink=11 hdist=1 tpe tbo qtrim=rl trimq=10 maq=10”. The quality-filtered reads were aligned with the genome of *A. tenuis* by HISAT2 ^37^, followed by determination of coral expression profiles by StringTie ^38^, as explained by Pertea et al ^39^. Unmapped reads were assembled using rnaSPAdes ^40^, and contigs derived from zooxanthellae were identified by similarity search against the genome sequences of corals and zooxanthellae using BLASTn. The unmapped reads were aligned to the contigs to calculate zooxanthella expression profiles using RSEM ^41^.

### Microbiome profiling by 16S rRNA amplicon sequencing

The V1–V2 region of 16S rRNA was amplified by RT-PCR using the primers PGM- 27F (CCATCTCATCCCTGCGTGTCTCCGACTCAG-[MID]- GATAGAGTTTGATCMTGGCTCAG) and PGM-338R (CCTCTCTATGGGCAGTCGGTGATTGCTGCCTCCCGTAGGAGT) and sequenced using IonPGM (Thermo Fisher Scientific, Waltham, MA, USA) to profile bacterial community structure. The sequence reads were preprocessed as follows: (i) the regions corresponding to PCR primers were trimmed using TagCleaner ^42^; (ii) the reads expected to contain >1% errors were removed with USEARCH ^43^; and (iii) the reads shorter than 270 bp were excluded. The preprocessed reads were truncated to 270 bp and clustered into OTUs using UPARSE ^44^. OTUs were taxonomically classified using RDP Classifier ^45^, and were designated as “undetermined” if their bootstrap values were below 80. A phylogenetic tree for the OTUs was constructed using FastTree ^46^ after multiple alignments with MAFFT ^47^. UniFrac analysis, based on the constructed tree, was performed using the R package phyloseq ^48^.

### Determination of zooxanthella cell density

To calculate zooxanthella cell density in coral branches (cells per square centimeter), the number of zooxanthella cells in coral tissue homogenates and the surface area of the corresponding coral skeleton with the tissue completely removed were determined. Homogenates of *A. tenuis* branches were prepared using AirFloss (Philips, Amsterdam, Netherlands) filled with Mg^2+^- and Ca^2+^-free artificial seawater (NaCl, 401.8 mM; Na_2_SO_4_, 27.6 mM; Na_2_HCO_3_, 2.29 mM; KCl, 8.91 mM; KBr, 0.81 mM; NaF, 0.62 mM; H_3_BO_4_, 0.39 mM; SrCl_2_, 11.3 μM; EDTA-Na_2_, 20 mM). The homogenate volume was determined, and zooxanthella cells were counted in a small portion of the homogenate using a 20 μm-depth hemocytometer. Surface area of the coral skeleton was measured as described previously ^49^.

### Genome assembly of *Endozoicomonas* isolate

Coral branches were rinsed in sterilized PBSE (phosphate-buffered saline with 20 mM EDTA). The tissue was then picked off using a sterilized toothpick, plated on marine broth supplemented with cycloheximide (100 g/mL), and incubated at 27 °C for 2 weeks. The obtained bacterial isolate was identified as *Endozoicomonas* species OTU5 by Sanger sequencing of 16S rRNA gene, and was grown in marine broth at 27 °C for additional 2 weeks. Genomic DNA was extracted by using the DNeasy Plant Mini Kit (QIAGEN), and pair-end libraries were prepared using the Nextera Library Prep Kit (Illumina) and sequenced using the Illumina MiSeq platform (Illumina). Low-quality reads were removed, and sequence adaptors were trimmed using fastp ^50^. Genome assembly of the quality-filtered sequence reads was conducted using SPAdes ^51^ in “careful” mode.

### Genome analysis of *Endozoicomonas* isolate

Publicly available genome sequences of 10 *Endozoicomonas* strains were downloaded from RefSeq (ftp://ftp.ncbi.nlm.nih.gov/genomes/refseq/bacteria/) for comparative genome analysis. Gene regions in each genome were predicted by using Prokka ^52^, and their functions were annotated using KAAS ^53^. Similar genes were grouped into orthogroups using Orthofinder ^54^. The genomes were aligned by using progressiveMauve ^55^ to define locally collinear blocks (LCBs), i.e., conserved genome regions among strains.

## Statistics

### Co-expression network analysis

Genetic modules in corals and zooxanthellae were determined based on weighted gene co-expression network analysis using the R package WGCNA ^45^. Genes expressed in <50% of the samples were excluded. The power parameter (coral: 14; zooxanthellae: 23) was selected to achieve good fitness (R^2^ > 0.8) for scale-free topology of the co-expression network.

### Gene set analysis

Functional annotation of the transcripts was performed using two methods. (i) Uniprot IDs were assigned to the transcripts by BLASTx search against the Swiss Prot database. UniProt IDs of the top hits were converted to gene ontology (GO) terms. (ii) KEGG orthology (KO) IDs were assigned to the gene clusters using KAAS ^53^. Gene set analysis was performed by Fisher’s exact test using the R package ClusterProfiler ^56^.

### Data availability

Raw RNA-seq reads and 16S rRNA amplicon sequences were deposited in NCBI under the BioProject accession numbers PRJNA743235 and PRJNA742893. The genome sequence of *Endozoicomonas* sp. ISHI1 is available at NCBI Genome under the accession number JAGRPU000000000.

### Code availability

The source codes and details of the analyses performed in this paper are available at https://tmaruy.github.io/coral_multi_omics.

## Acknowledgements

This work was supported by JST-CREST (grant number JPMJCR12A4). The supercomputing resource was provided by the Human Genome Center (the University of Tokyo, Tokyo, Japan).

## Author contributions

TM analyzed data. MI, SW, YO, TM and YNi collected the samples and sequenced them. MI, YO, YNi, and KI isolated the bacteria and sequenced the bacterial genome. HF, SS, and YNa performed environmental monitoring. HT, CS, NS, and KY guided the research. TM and HT wrote the manuscript.

## Materials and Correspondence

Correspondence to Haruko Takeyama.

## Tables

**Table S1** Number of collected samples

**Table S2** Proportion of RNA-seq reads mapped onto A. tenuis genome

**Table S3** Gene set analysis for coral gene modules with Gene Ontology

**Table S4** Gene set analysis for coral gene modules with KEGG Pathway

**Table S5** Gene set analysis for zooxanthellae gene modules with Gene Ontology

**Table S6** Gene set analysis for zooxanthellae gene modules with KEGG Pathway

**Table S7** Differentially expressed genes in corals harboring Anaplasma bacteria

**Table S8** Differentially expressed genes in corals harboring Endozoicomonas bacteria

**Table S9** Gene set analysis for differentially expressed genes in corals harboring Endozoicomonas

**Table S10** Comparative genome analysis of Endozoicomonas species

---

**Supplementary Fig. 1.**
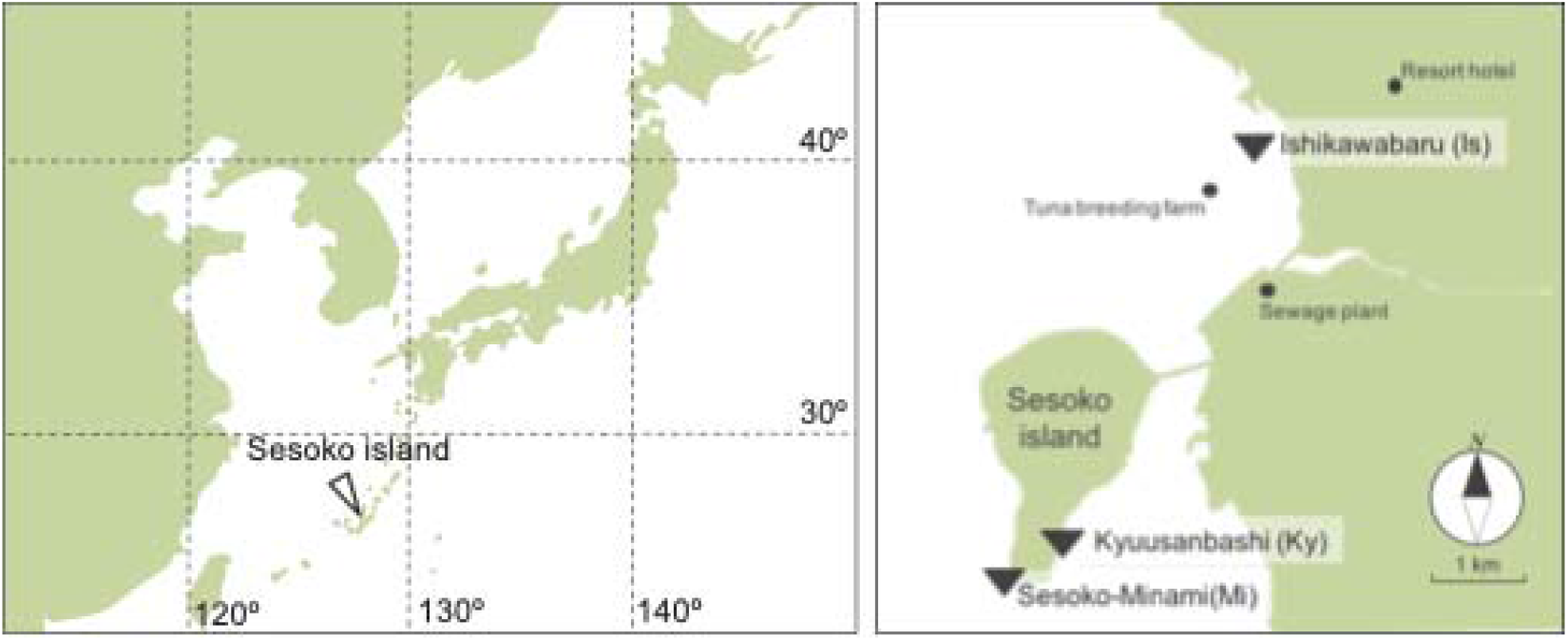
Sampling site. Scleractinian corals *Acropora tenuis* were sampled at the three sampling sites (*Is*, *Ky*, *Mi*) around Sesoko island, Okinawa, Japan.

**Supplementary Fig. 2.**
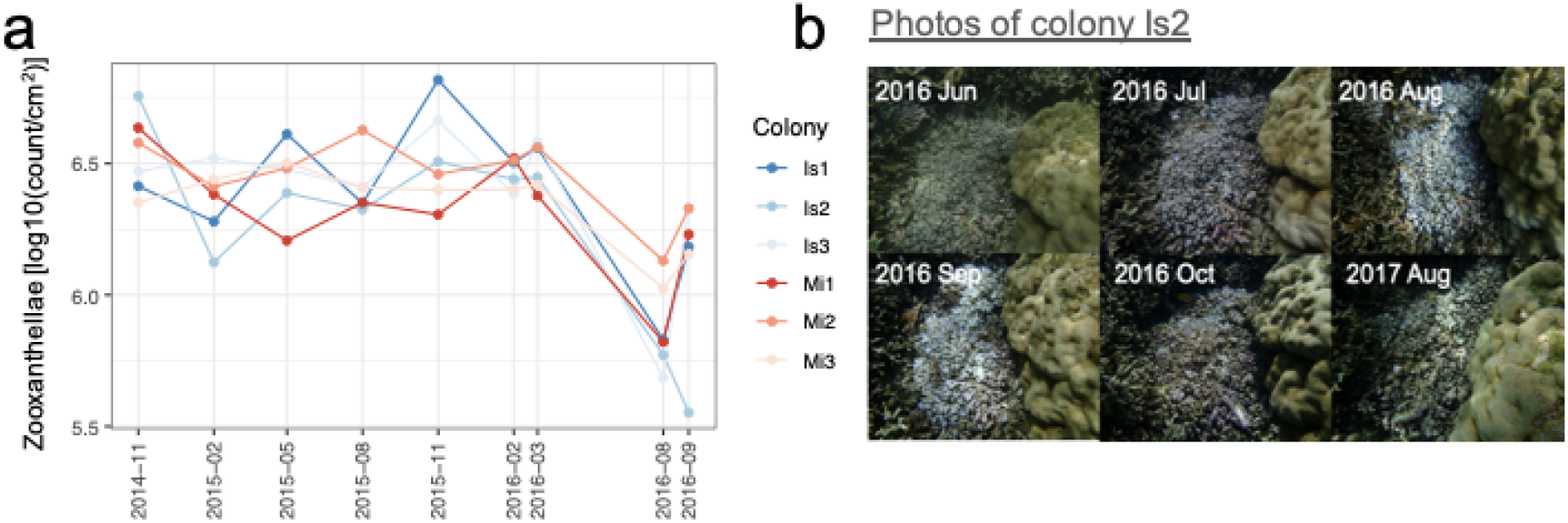
Changes in zooxanthellae densities over time. **a** Seasonal change of zooxanthellae densities in each coral colony. **b** Representative photos of Is2, a colony which showed prominent bleaching in 2016 Aug and 2016 Sep.

**Supplementary Fig. 3.**
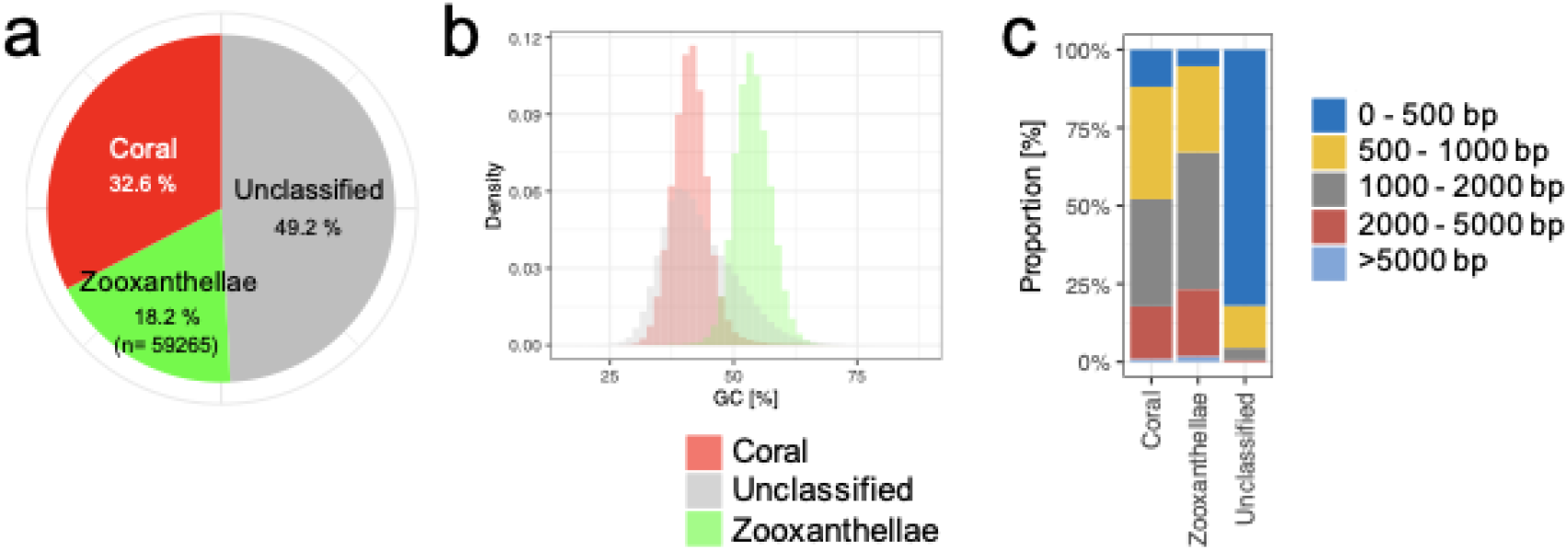
*De novo* transcriptome assembly using unmapped reads. **a** Proportion of the contigs derived from coral and zooxanthellae. Contigs of coral and zooxanthellae were classified by similarity search against transcripts of *A. tenuis* and zooxanthellae. Contigs showing no similarity with both transcripts were represented as unclassified. **b** GC contents of the contigs. **c** Length of the contigs.

**Supplementary Fig. 4.**
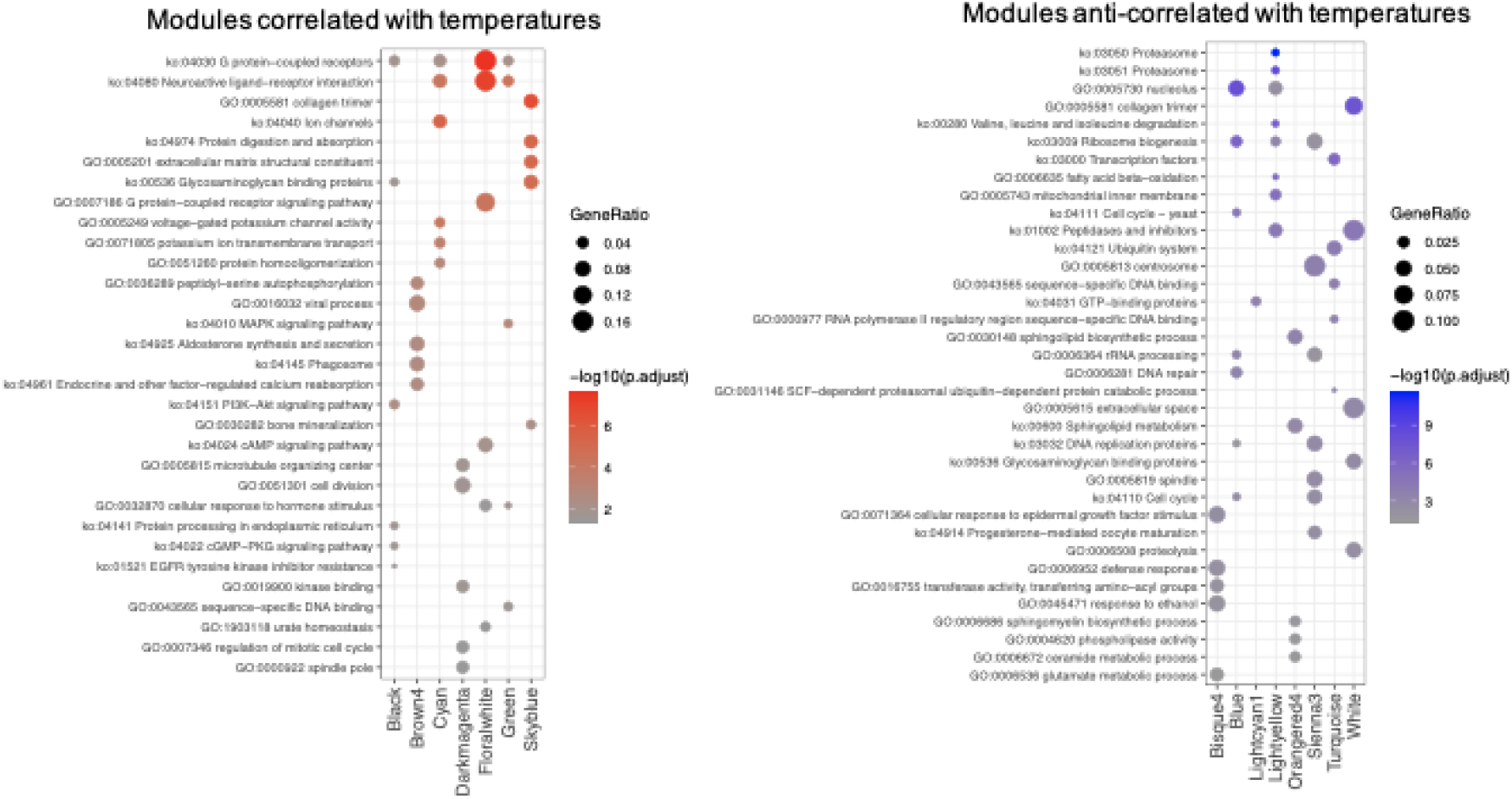
Function of temperature-dependent modules in corals. Dot plots depict gene ontology (GO) terms and KEGG pathways significantly associated with the temperature-dependent modules. Colors and sizes of the dots correspond to adjusted p-values and proportion of genes belonging to the functional groups in the modules. Statistical significance was calculated by Fisher’s exact test.

**Supplementary Fig. 5.**
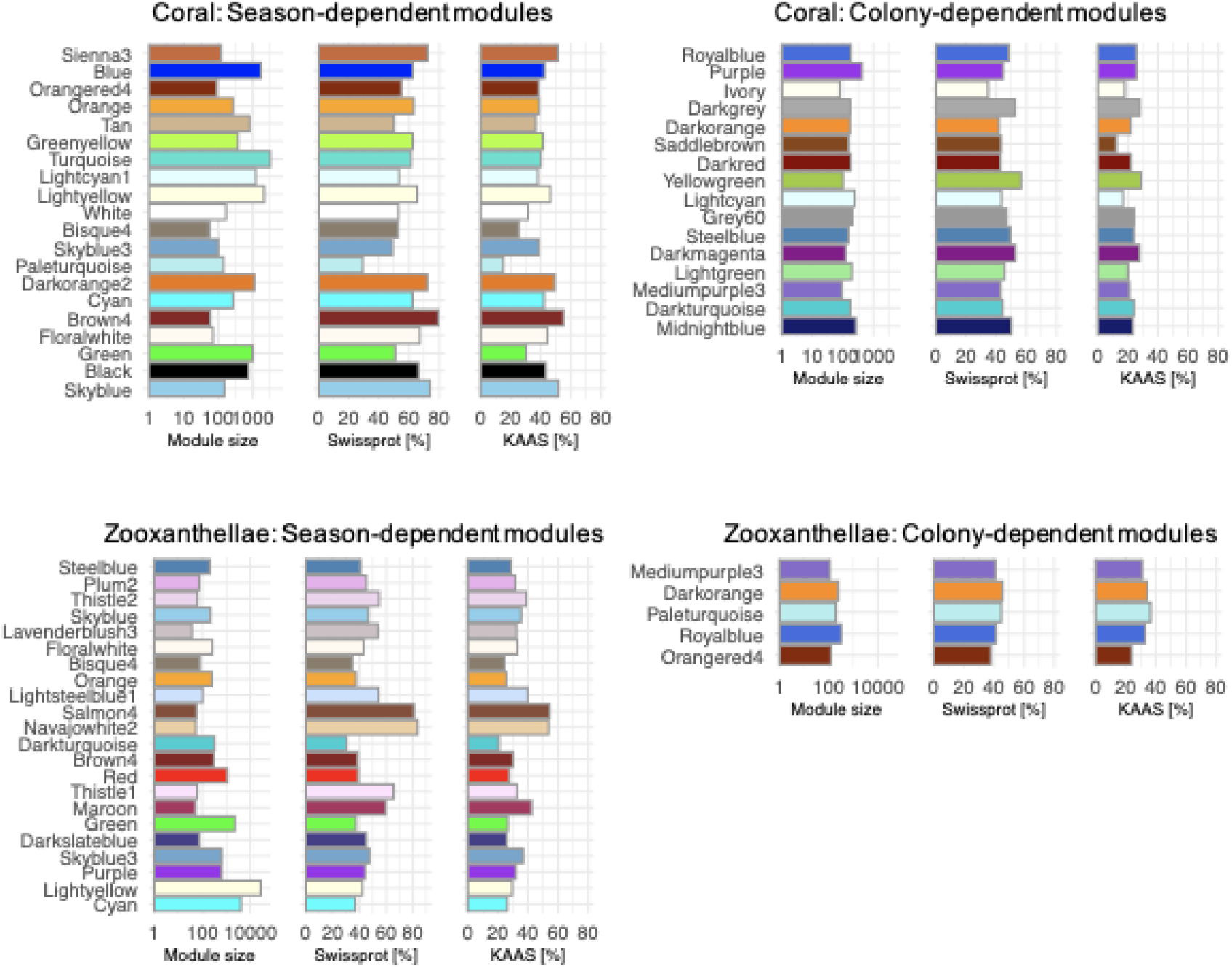
Size and annotation rate of the modules. Bar plots represent module size, proportion of genes showing similarities with sequences in Swissprot and KEGG.

**Supplementary Fig. 6.**
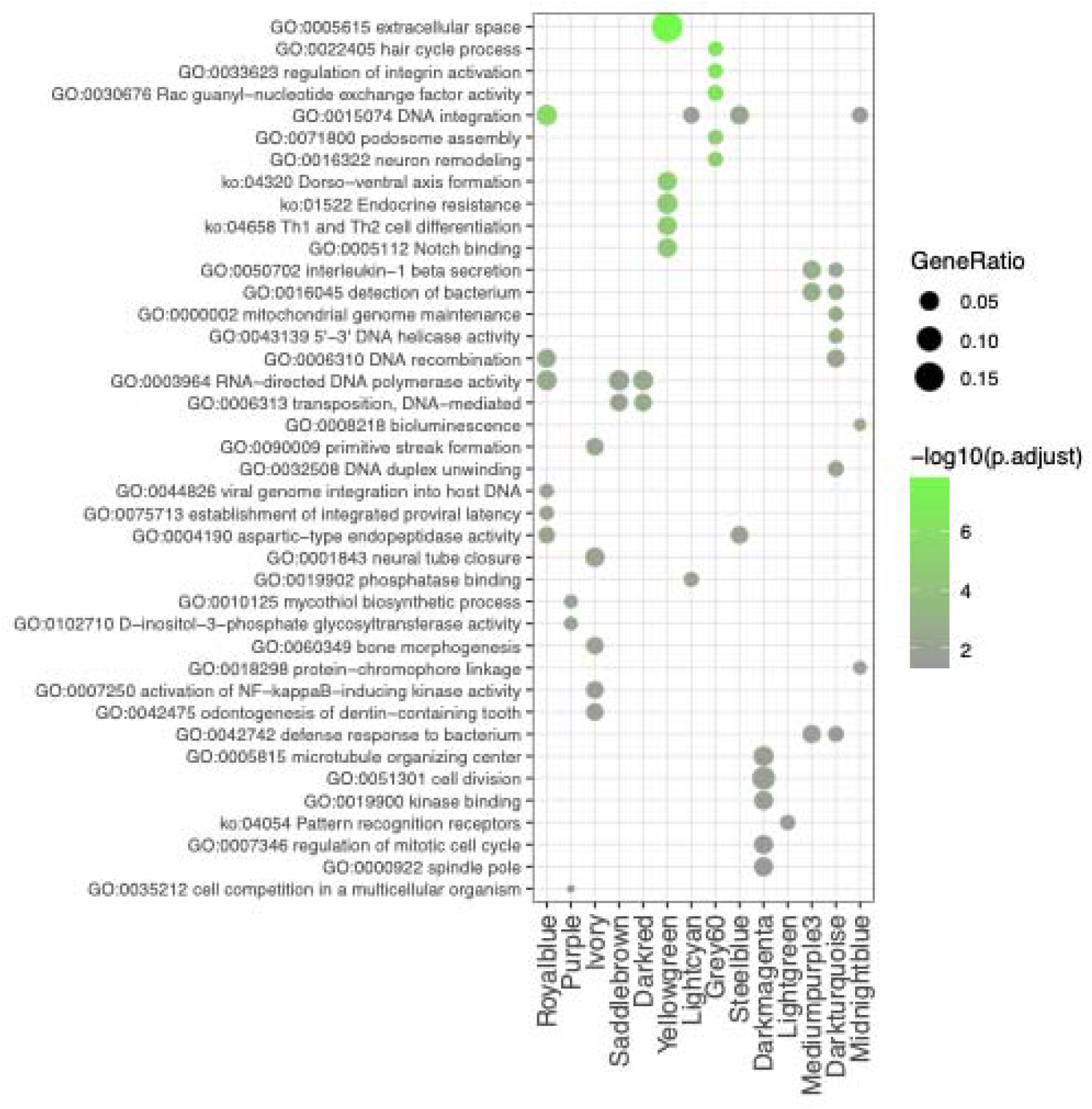
Function of colony-dependent modules in corals. Dot plots depict gene ontology (GO) terms and KEGG pathways significantly associated with the colony-dependent modules. Colors and sizes of the dots correspond to adjusted p-values and proportion of genes belonging to the functional groups in the modules. Statistical significance was calculated by Fisher’s exact test.

**Supplementary Fig. 7.**
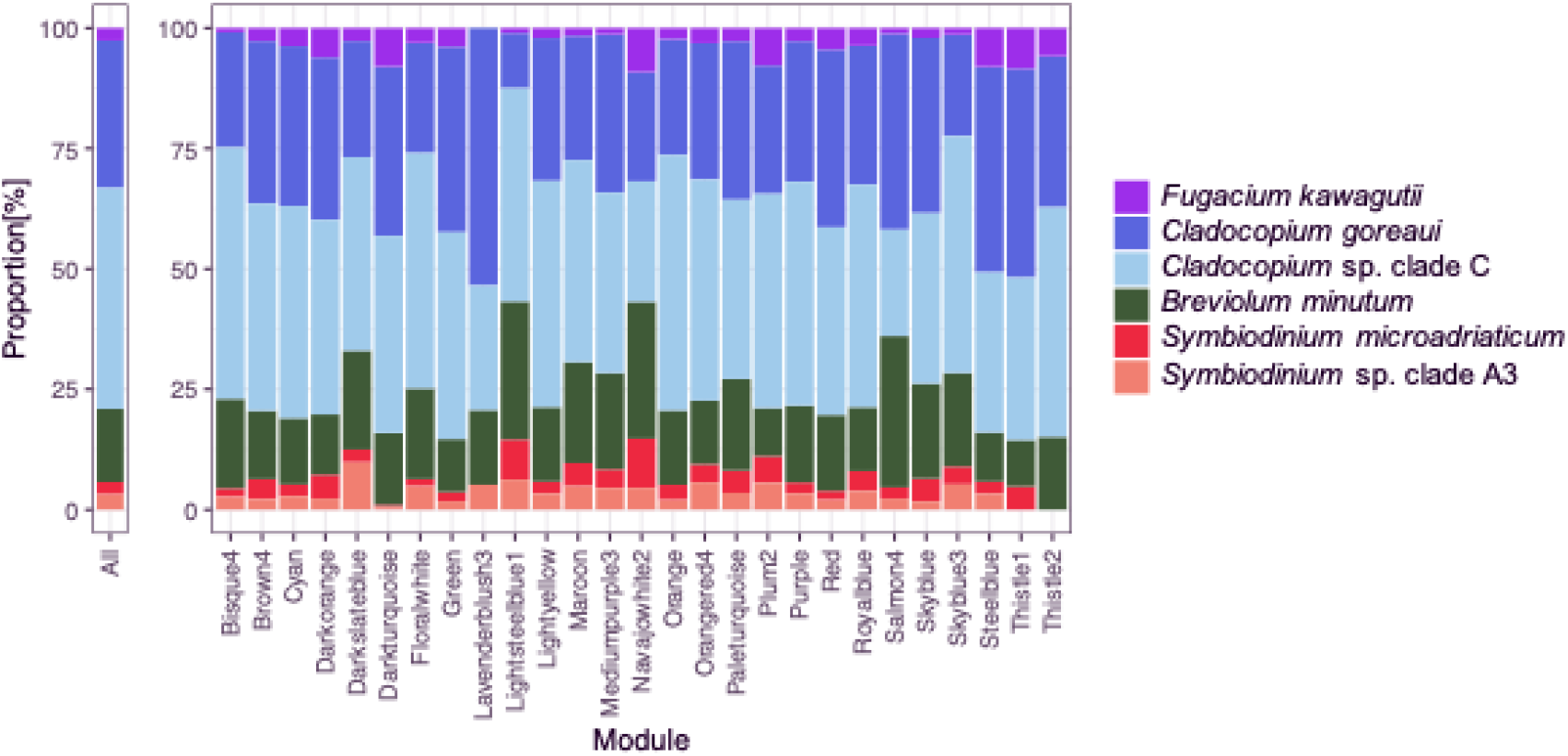
Taxonomic assignment for zooxanthellae transcripts. Bar chart depicts the taxonomic composition of zooxanthellae transcripts. Similarity searches against protein sequences of six zooxanthellae species were conducted. Each transcript was classified based on the origin of the best hit sequence. *Symbiodinium* sp. clade A3, *S. microadriaticum* (clade A), *Breviolum minutum* (clade B), *Cladocopium* sp. clade C, *C. goreaui* (clade C1) and *Fugacium kawagutii* (clade F).

**Supplementary Fig. 8.**
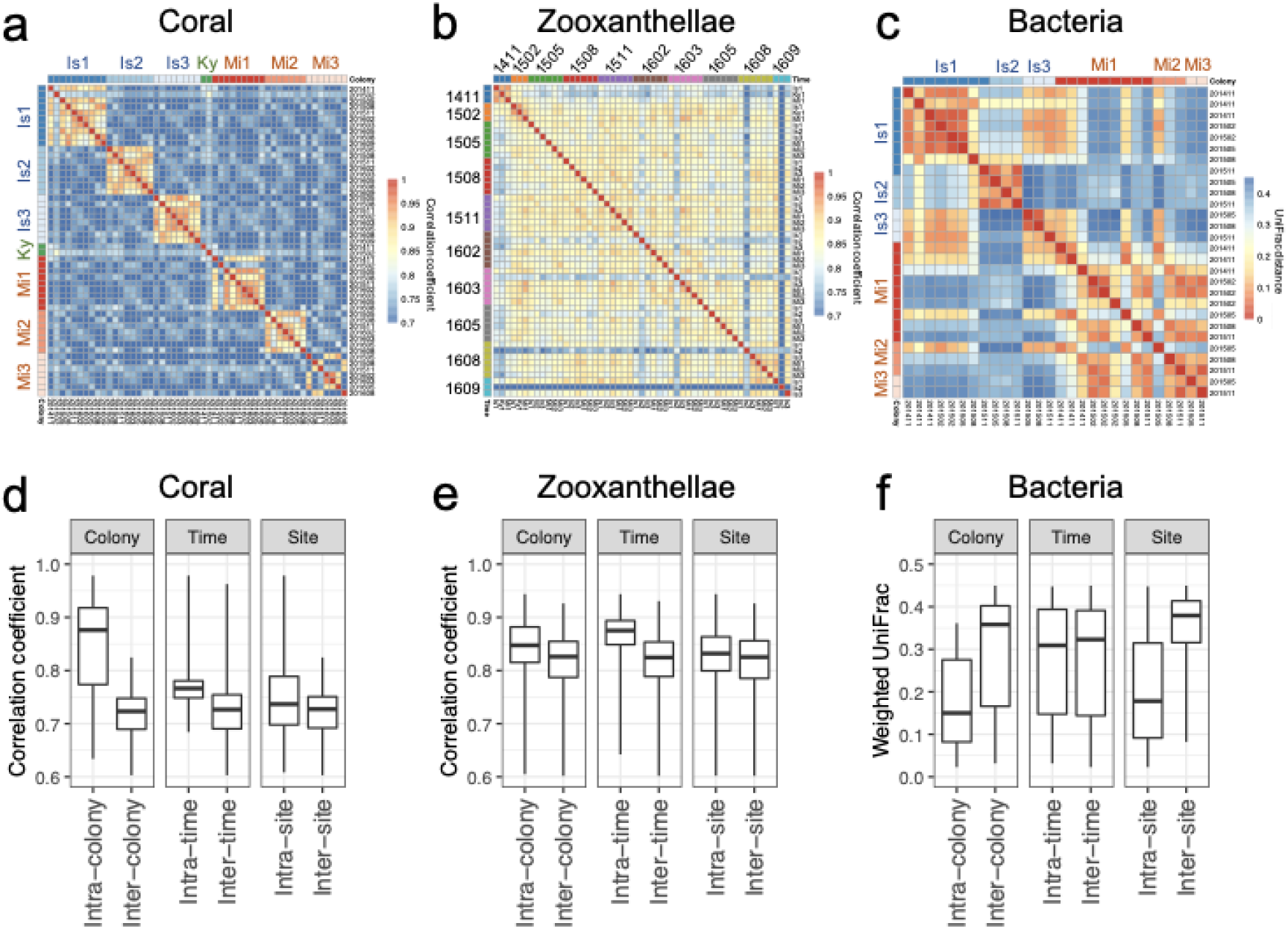
Similarities of coral transcriptome, zooxanthellae transcriptome and microbiome among samples. **a–c** Similarities of **a** coral transcriptome, **b** zooxanthellae transcriptome and **c** microbiome among samples. Pearson’s correlation coefficient was used as a similarity indicator for transcriptome of coral and zooxanthellae. Weighted UniFrac distance was used for microbiomes. **d–f** Box plot of the similarities. Midlines in the box represent medians. Upper and lower hinges represent 25 % quantiles and 75 % quantiles. Upper and lower whiskers represent maximum and minimum values.

